# N-glycosylation of the protein disulfide isomerase Pdi1 ensures *Ustilago maydis* virulence

**DOI:** 10.1101/571125

**Authors:** Miriam Marín-Menguiano, Ismael Moreno-Sánchez, Ramón R. Barrales, Alfonso Fernández-Álvarez, José Ignacio Ibeas

## Abstract

Fungal pathogenesis depends on accurate secretion and location of virulence factors which drive host colonization. Protein glycosylation is a common posttranslational modification of cell wall components and other secreted factors, typically required for correct protein localization, secretion and function. Thus, the absence of glycosylation is associated with animal and plant pathogen avirulence. While the relevance of protein glycosylation for pathogenesis has been well established, the main glycoproteins responsible for the loss of virulence observed in glycosylation-defective fungi have not been identified. Here, we devise a proteomics approach to identify such proteins and use it to demonstrate a role for the highly conserved protein disulfide isomerase Pdi1 in virulence. We show that efficient Pdi1 N-glycosylation, which promotes folding into the correct protein conformation, is required for full pathogenic development of the corn smut fungus *Ustilago maydis*. Remarkably, the observed virulence defects are reminiscent of those seen in glycosylation-defective cells suggesting that the N-glycosylation of Pdi1 is necessary for the full secretion of virulence factors. All these observations, together with the fact that Pdi1 protein and RNA expression levels rise upon virulence program induction, suggest that Pdi1 glycosylation is a crucial event for pathogenic development in *U. maydis*. Our results provide new insights into the role of glycosylation in fungal pathogenesis.

**Author summary:** Fungal pathogens require virulence factors to be properly secreted and localized to guarantee complete infection. In common with many proteins, virulence factors must be post-translationally modified by glycosylation for normal localization, secretion and function. This is especially important for virulence factors, which are mainly comprised of cell wall and secreted proteins. Aberrant glycosylation leads to a loss of virulence in both animal and plant pathogenic fungi. We have previously demonstrated that glycosylation is important for virulence of the corn smut fungus, *Ustilago maydis*. However, the glycoproteins involved and their specific roles in the infection process have not yet been reported. Here, we describe a proteomic assay designed to identify glycoproteins involved in plant infection. Using this method, we define the role of Pdi1 protein disulfide isomerase in virulence. Interestingly, abolishing Pdi1 N-glycosylation mimics Δ*pdi1* defects observed during infection, suggesting that Pdi1 N-glycosylation is required for the secretion of virulence factors. We hypothesize that Pdi1 N-glycosylation is crucial for maintaining proper effector protein folding during the infection process, especially in the harsh conditions found inside the maize plant.

## Introduction

Protein glycosylation is a common eukaryotic post-translational mechanism required for the correct folding, activity and secretion of many proteins. Glycosylation involves the synthesis and addition of different polysaccharide cores (sugars) to specific amino acids within a consensus sequence. Most glycoproteins are plasma membrane-associated cell wall and secreted proteins, which acquire glycosyl groups during their transit through the Endoplasmic Reticulum (ER) and Golgi Apparatus (GA) (1,2). Defects during the synthesis or addition of sugars to target proteins affect many biological processes; for instance, impaired human protein glycosylation causes more than 100 severe embryonic development disorders (3). In pathogenic fungi, glycosylation defects lead to a reduction or absence of virulence in plant and animal pathogens (4–8).

Protein glycosylation is divided into different types based on the structure and composition of the oligosaccharide cores and the amino acids to which they are attached. N- and O-glycosylation are the most common types in pathogenic fungi. N-glycosylation consists of the addition of an oligosaccharide core, composed of two N-acetylglucosamines (NAcGlc), nine mannoses (Man) and three glucose (Glc) molecules, NAcGlc_2_Man_9_Glc_3_, to the nitrogen chain of an asparagine residue in the sequence Asn-*x*-Ser/Thr, where *x* can be any amino acid except proline (9,10). O-glycosylation is more variable than N-glycosylation in terms of the types of sugars added. In fungi O-mannosylation is the most common type of O-glycosylation and is characterized by the addition of Man residues to target proteins. In contrast to N-glycosylation, O-glycosylation involves sequential additions of Man to the oxygen chain of Ser or Thr amino acids although no amino acid consensus sequence has been identified (11). N- and O-linked glycans are later processed during their transit across the ER and GA, and specific trimming of sugars is also essential for the function and secretion of glycoproteins (5,12).

Crucial components for fungal pathogenesis belonging to N- and O-glycosylation pathways have been identified in several organisms such as *Candida albicans, Aspergillus nidulans, Cryptococcus neoformans, Magnaporthe oryzae* or *Ustilago maydis* (4,6–8,13–15). The loss of these proteins primarily affects those stages of pathogenic development that require robust glycoprotein secretion. The involvement of protein glycosylation in fungal virulence has been extensively explored in the corn smut fungus *U. maydis* (4,5,16).

*U. maydis* combines both pathogenic and non-pathogenic life cycles. During the non-pathogenic cycle, *Ustilago* grows as haploid yeast-like cells that can be easily cultured in the laboratory. The pathogenic cycle starts when two sexually compatible strains mate on the maize plant surface. Sexual compatibility is determined by two independent loci: *locus a*, which encodes a pheromone-receptor system; and *locus b*, which encodes a transcription factor formed by a *bE/bW* heterodimer. Formation of the *bE/bW* complex triggers development of an infective dikaryon filament (17). Physical and chemical plant signals are sensed by filaments that develop a morphogenetic structure called the appressorium, which mediates plant cuticle penetration (18). Once inside plant tissues, the fungus expands in a branched filamentous form generating hypertrophied plant cells, macroscopically visible as tumors (19–23).

N- and O-glycosylation are both crucial for *U. maydis* pathogenic development. The loss of the O-mannosytransferase Pmt4, which catalyzes the addition of mannoses to target proteins, compromises both appressorium formation and plant cuticle penetration. Hence, Δ*pmt4* cells are unable to invade the plant tissues and tumor induction is fully abolished (4). Glucosidase I and II (Gls1 and Gas1/Gas2) are important for N-glycan processing in the ER and play crucial roles during the early stages of *U. maydis* plant colonization (5,8). A reasonable explanation for these drastic virulence defects is that deficient glycosylation could greatly alter the location and/or function of glycoproteins involved in virulence, and consequently compromise multiple stages of *U. maydis* pathogenic development such as plant cuticle penetration, fungal progression inside plant tissues or plant defense responses. Despite the importance of Pmt4 and Gls1 for maize infection, the virulence factors glycosylated by these proteins are still poorly described. In this context, an *in silico* search for putative Pmt4 targets identified Msb2 as an O-glycoprotein, whose deletion causes virulence defects that resemble some of those described for the Δ*pmt4* mutant (16). However, a more wide-ranging approach is required to find new glycosylated virulence factors.

In this work we devise a proteomics approach designed to identify N- and O-glycoproteins produced when the infection program is activated. Using this method, we identify several Gls1 and Pmt4 targets involved in virulence. Among these, we further characterize Pdi1, a disulfide isomerase protein whose glycosylation we demonstrate to be required for full virulence in maize plants due to its involvement in glycoprotein folding. Furthermore, we show that the deletion of Pdi1 affects glycoprotein secretion in *U. maydis*. We speculate about its role during the infection process, which could be related to ensuring the effective production and secretion of many virulence factors.

## Results

### Glycoproteomic screening for Pmt4 and Gls1 substrates

Previous work from our laboratory have shown that the *U. maydis* proteome contains a high number of putative O-glycoproteins mannosylated by the O-mannosyltransferase Pmt4 (16). An *in silico* screen for proteins harboring Ser/Thr-rich regions, where Pmt4 attaches mannoses, revealed that around 65% of *U. maydis* proteins are potential O-glycosylation targets (16). If proteins containing N-glycosylation sites are included, this number rises to over 70% of the proteome, suggesting a potential role for protein glycosylation in the activity of a high proportion of *U. maydis* proteins (Pérez-Pulido, personal communication). In order to identify virulence-related glycosylation targets, we designed a selective glycoproteomic screen based on the hypothesis that glycoproteins whose expression is modified upon virulence program induction are likely to have important roles in the pathogenic phase of the *U. maydis* life cycle.

For this screen we set out to analyze cell extracts corresponding to cytosolic, cell wall and secreted proteins using two-dimensional differential gel electrophoresis (2D-DIGE) to detect glycoproteins whose spot area or intensity were altered by the loss of Pmt4 or Gls1 and corresponding effect on O- or N-glycosylation activities, respectively. To isolate glycoproteins from cytosolic extract and avoid interference with other proteins, total protein extracts were enriched for mannose-containing proteins by High-Performance Liquid Chromatography (HPLC) using *concanavalin A* columns (Fig 1). Due to the high proportion of glycoproteins in cell wall and secreted extracts, HPLC enrichment was not applied to these preparations. Entry into the virulence phase was controlled by expressing Biz1, a *b*-dependent zinc-finger protein whose induction activates pathogenic filamentous growth, under the control of the carbon source-regulated *P*_*crg*_ promoter that is induced by arabinose (*biz ON*) and repressed by glucose (*biz OFF*) (24,25). Thus, wild-type (*wt*), Δ*pmt4* and Δ*gls1* cells harboring *P*_*crg*_ controlled-*biz1* were collected under inducing and repressing conditions and protein samples compared, using three replicas of each, to identify the differentially migrating proteins (Fig 1).

**Fig 1.**
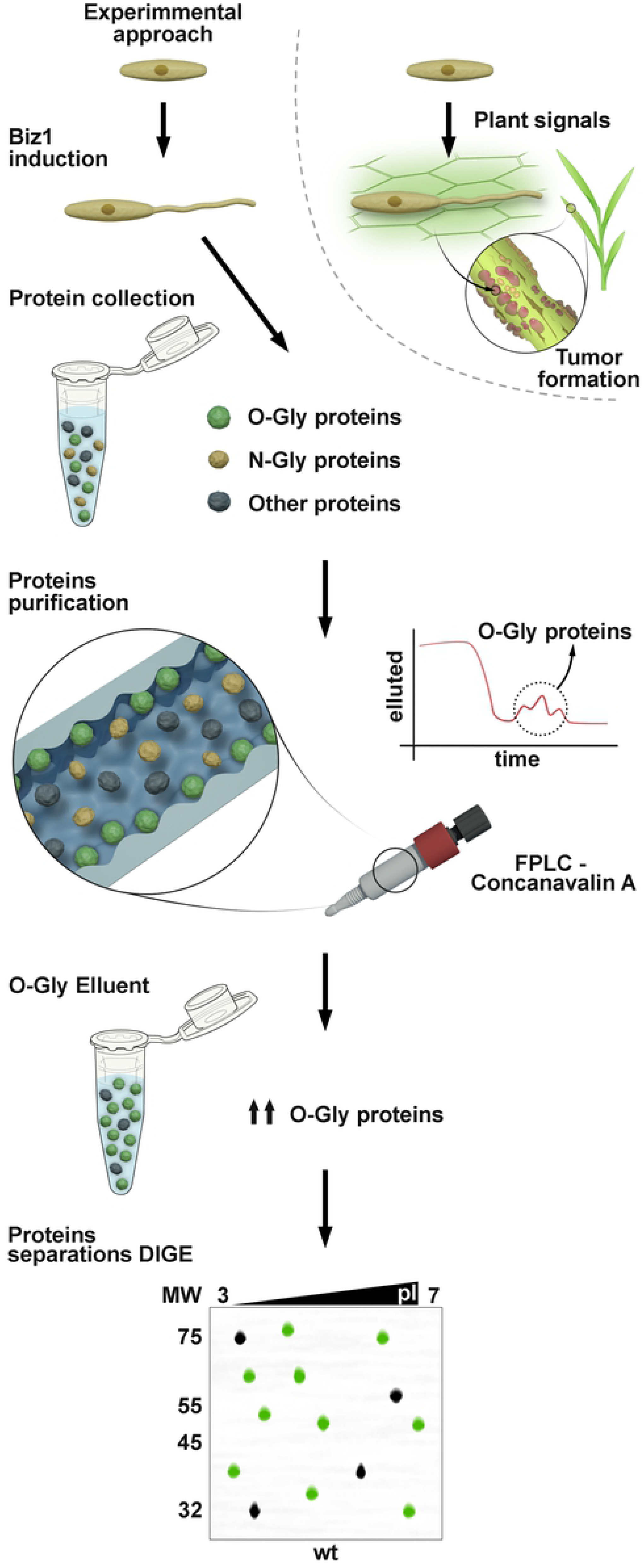
Proteomic approach to identify glycoproteins involved in virulence. *Biz1* overexpression is used to induce the pathogenic pathway and cytosolic proteins are then collected from wild-type *U. maydis* and from a mutant deficient in either O- or N-glycosylation. O- or N-glycoproteins are purified through a Concanavalin A FPLC column using O- or N-glycoprotein specific eluents. Finally, glycoproteins are tagged with fluorophores and separated by 2D electrophoresis.

By applying the approach described above, we detected protein spots whose areas showed altered electrophoretic mobility compared to *wt* in a Biz1-dependent manner in both Δ*pmt4* and Δ*gls1* cells, presumably corresponding to O- and N-glycoproteins, respectively. Using MALDI–MS we identified four proteins in the cytosolic extract with altered mobility in both mutants that probably correspond to both O- and N-glycosylated proteins: the chorismate mutase Cmu1, the α-L-arabinofuranosidase Afg1, the invertase Suc2 and Um10156, a putative disulfide-isomerase (Fig 2). In addition, we identified 27 N- or O-glycoproteins from the cytosolic extract, 6 from the cell wall extract and 11 from the secreted extract showing electrophoretic mobility changes (Fig 3, S1 Table). Of the proteins identified only Cmu1, Um04926 (Pep4), Afg1 and Um04309 (Afg3) have been previously characterized. Cmu1 is a secreted virulence factor that controls the metabolic status of plant cells during fungal colonization; deletion of *cmu1* reduces *U. maydis* virulence in the solopathogenic strain CL13 (26). Pep4 is a proteinase A located at the vacuole, which is involved in the yeast to micelium dimorphic transition and whose deletion causes reduced virulence including incomplete teliospore maturation (27). On the other hand, Afg1 and Afg3 are arabinofuranosidases that participate in the degradation of arabinoxylan, a plant cell wall component. While the loss of *afg1* or *afg3* is dispensable for virulence, the triple deletion of *afg1, afg2* and *afg3* compromises full pathogenic development (28). The involvement of the other identified proteins in pathogenesis remains to be determined.

**Fig 2.**
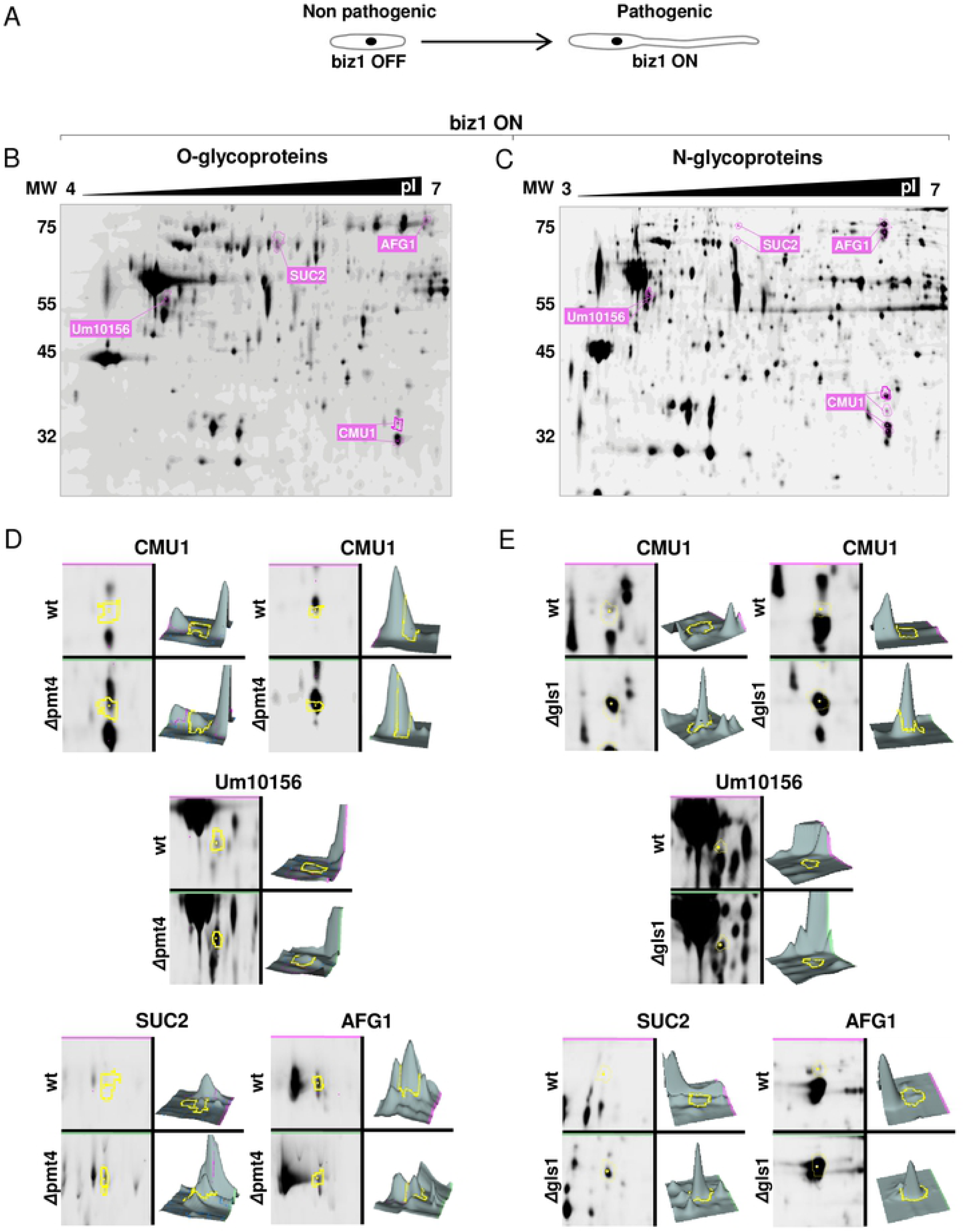
2D-DIGE expression analysis of Pmt4 and Gls1-dependent cytoplasmic glycoproteins. Cells growing in inducing conditions develop hyphae (A). Images of DIGE gels containing an internal standard loaded with equal amounts of each sample, showing protein changes between *biz1* and *pmt4*-mutants (B) or *biz1* and *gls1*-mutants (C). Protein expression profile changes between wild-type *biz1*^*crg*^ vs Δ*pmt4* mutants (D) or Δ*gls1* mutants (E) showed by the DeCyder software analysis. The edge of the indicated protein is displayed in the DIGE gel in pink and in a specific zoom in yellow.

**Fig 3.**
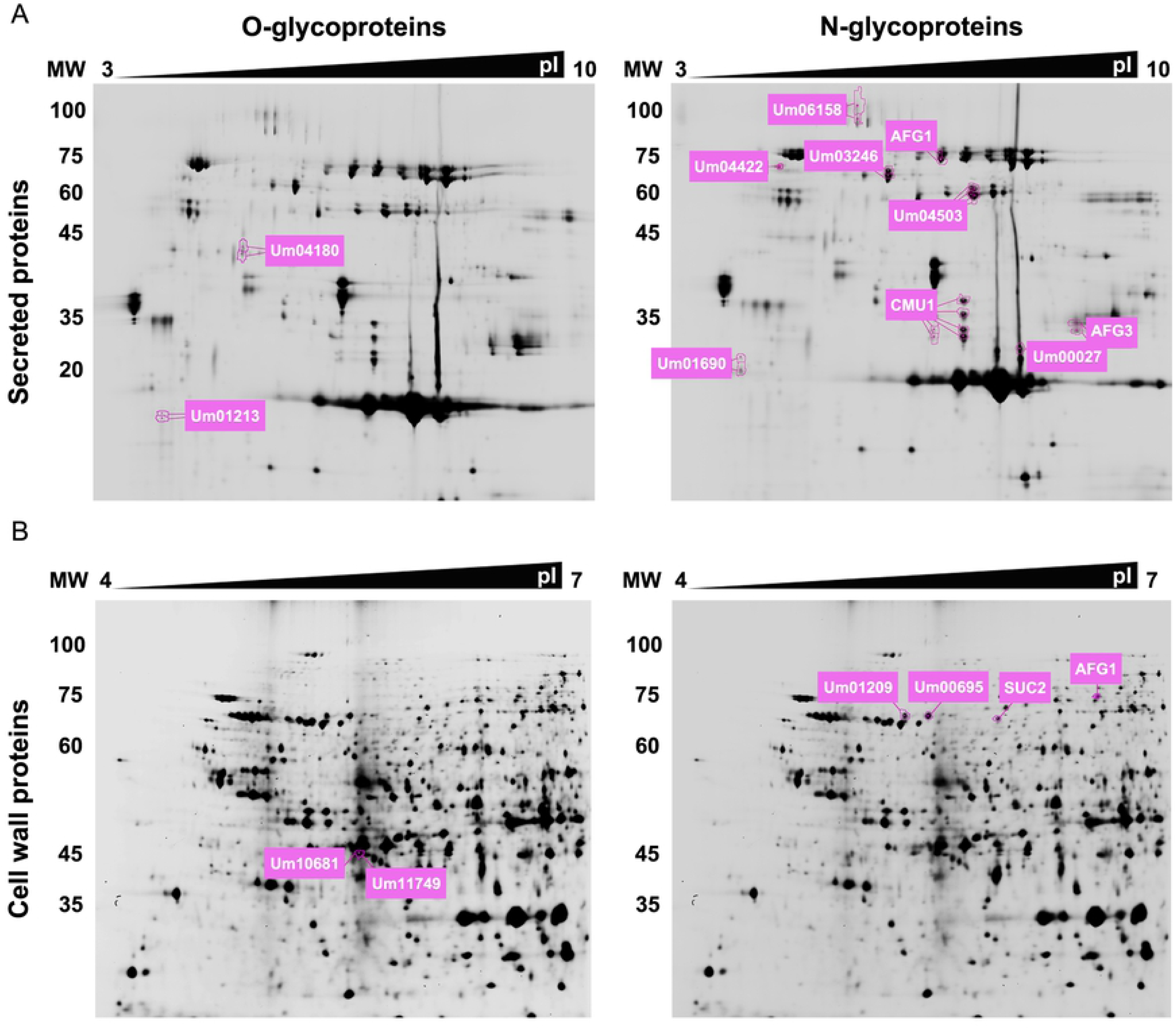
2D-DIGE expression analysis of Pmt4 and Gls1-dependent secreted or cell wall glycoproteins. Images of the DIGE gels containing an internal standard loaded with equal amounts of each samples, showing protein changes between wild-type *biz1*^*crg*^ and *pmt4*-mutants (secreted glycoproteins in A, left part; cell wall glycoproteins in B, left part) or wild-type *biz1*^*crg*^ and *gls1*-mutants (A, right part; B right part).

### The disulfide-isomerase Pdi1 is a substrate of Pmt4 and Gls1, and Pdi1 is required for pathogenesis

To determine the involvement in pathogenesis of the uncharacterized proteins, we compared the virulence capability of CL13 strain mutants carrying deletions corresponding to several of the candidate proteins. The CL13 strain harbors genes encoding a compatible *bE1/bW2* heterodimer allowing it to complete full pathogenic development (29). Analysis of disease symptoms after infection revealed strong virulence defects in Δ*afg1*, Δ*Um00309*, Δ*Um02751*, Δ*Um03416*, Δ*Um04180*, Δ*Um04422*, Δ*Um04733*, Δ*Um05223*, Δ*Um10156*, and Δ*Um11496* infections (Fig 4A and S1 Table). Candidate genes were then tested in more virulent strains; the wild type FB1 and FB2 strains (30), which have a higher infection capacity, and SG200, which carries the *mfa2* gene that encodes the compatible mating type pheromone (29) and confers stronger virulence. Based on the virulence assays in these backgrounds we identified Um10156 as having the most significant role in plant infection (Fig 4B, C and D). Blast analysis for Um10156 revealed 36% identity to *S. cerevisiae* Pdi1 with 48% and 43% identity in the two thioredoxin domains (Fig 5A and S1 Fig). Thus, we will now refer to Um10156 as Pdi1 for the remainder of the article. *S. cerevisiae* Pdi1 is a disulfide-isomerase protein, member of the thioredoxin superfamily of redox proteins whose canonical role is to support the folding of newly synthesized proteins in the ER *via* the addition of disulfide bonds (31).

**Fig 4.**
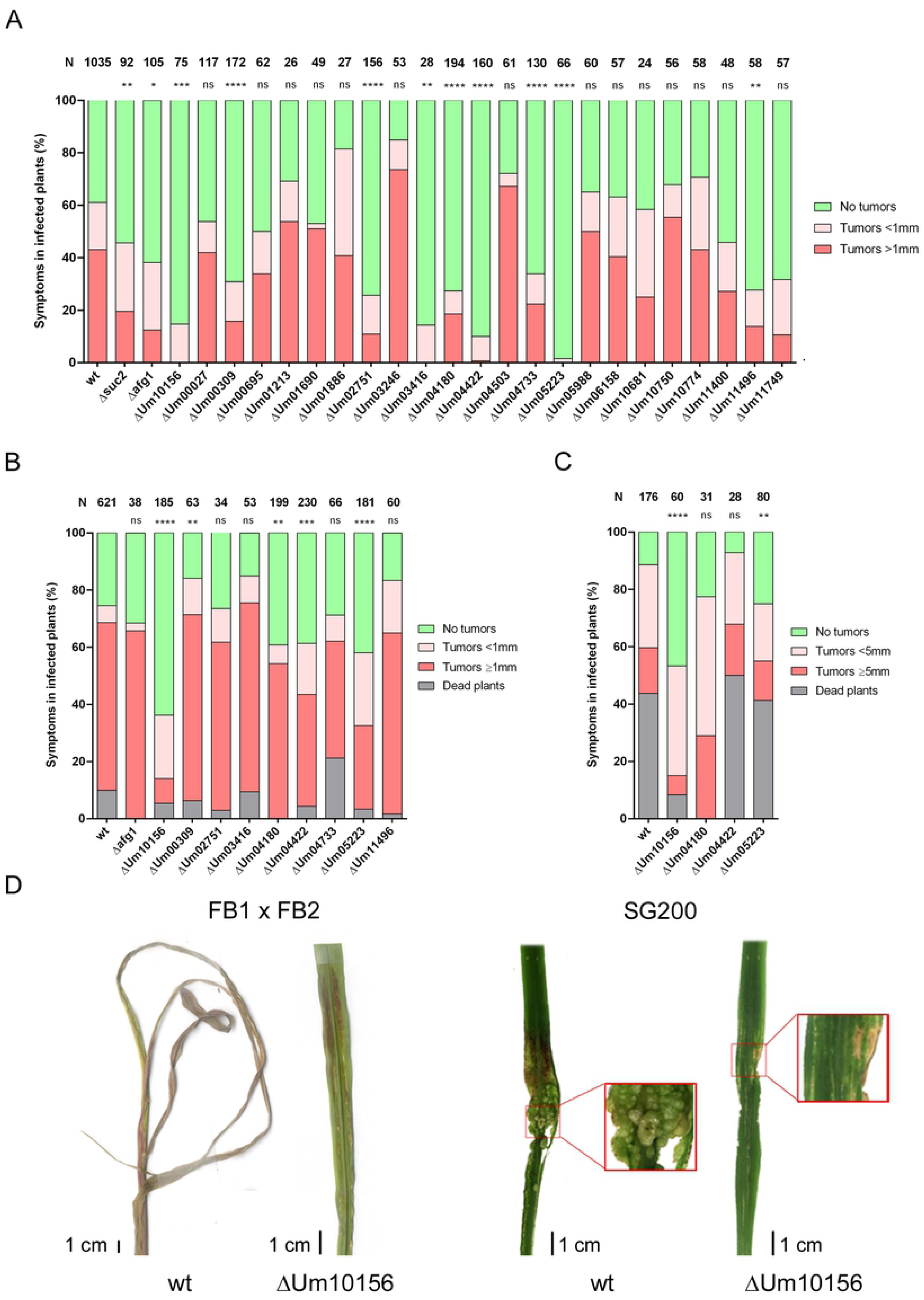
Infection assay of pathogenic pathway glycoprotein candidates. Deletions and infections were first carried out in the CL13 strain (A). Deletions showing a statistically significant reduction in virulence were then assayed in SG200 (B) and subsequently in FB1 and FB2 compatible strains (C). Total number of plants infected is indicated above each column. The Mann-Whitney statistical test was performed for each mutant versus the corresponding wild-type strain (ns: not statistically significant; * for p-value < 0.05; ** for p-value < 0.01; *** for p-value < 0.005; **** for p-value < 0.0001). (D) Representative Δ*Um10156* disease symptoms compared to wild-type.

**Fig 5.**
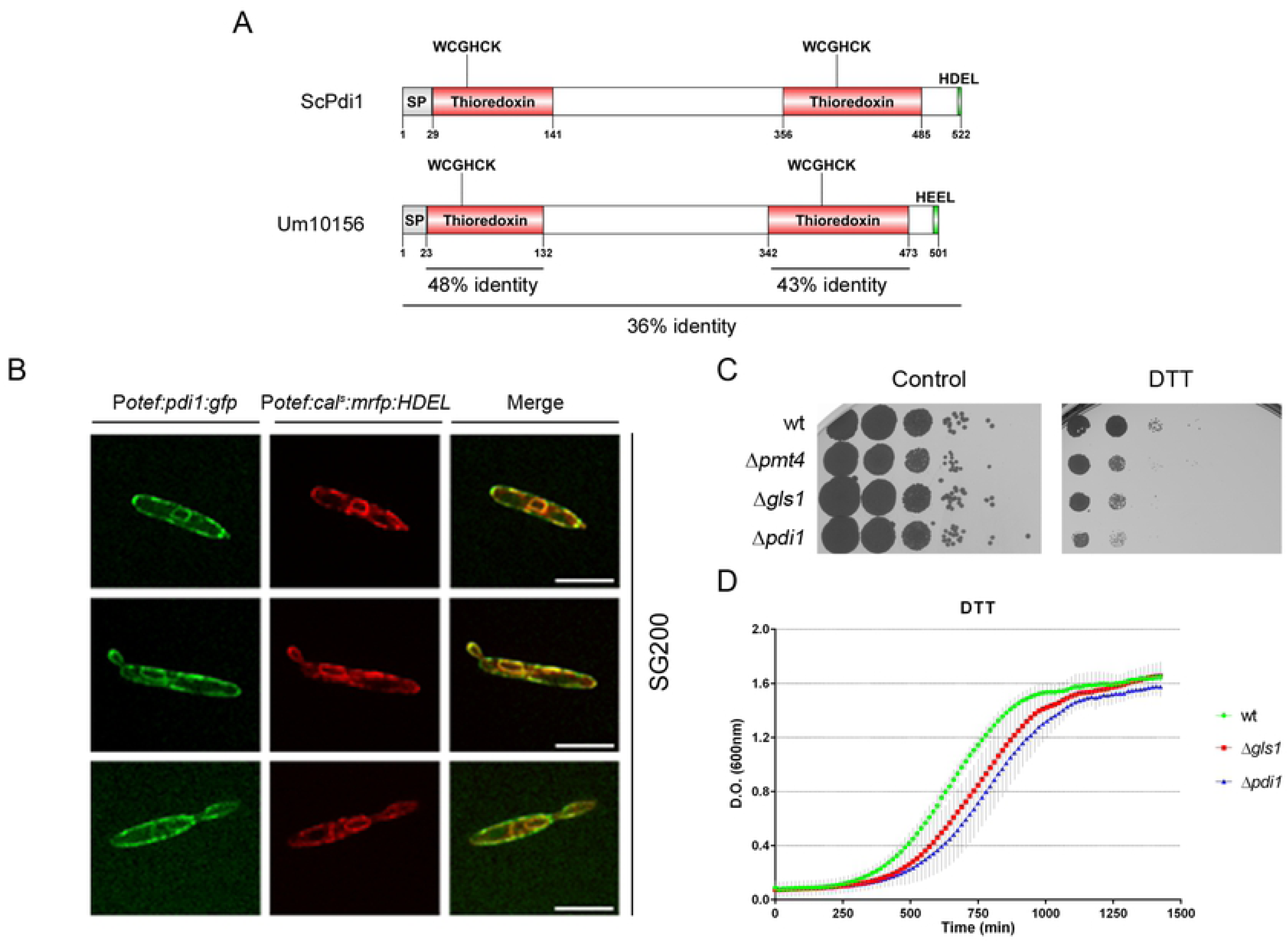
Pdi1 (Um10156) is a protein disulfide isomerase that localizes to the ER and its deletion results in sensitivity to ER stress inducing treatments. (A) Schematic representation of ScPdi1 and Um10156 showing signal peptides (SP), thioredoxin domains, WCGHCK active sites, HDEL/HEEL ER localization sites and thioredoxin domain and whole protein identity percentages obtained by BLASTp. (B) Pdi1:GFP co-localizes with the ER marker mRFP:HDEL. (C) ER stress assay was performed on solid CM plates supplemented with 2% D-glucose and 4 mM DTT. (D) ER stress assay performed with liquid media supplemented with 10 mM DTT. Scale bar represents 5 µm.

### The loss of Pdi1 compromises fungal expansion inside plant tissues

Previous studies in budding yeast have shown that Pdi1 function is critical under ER stress conditions, where a large number of misfolded proteins are produced (32,33). To confirm that this function of Pdi1 is retained in *U. maydis*, we explored Pdi1 localization and studied the ability of Δ*pdi1* cells to grow under reductive ER stress induced by DTT. Microscopy co-localization analysis of Pdi1:GFP and the ER marker mRFP:HDEL confirmed the localization of Pdi1 to this organelle (Fig 5B), where it has been shown to work in conjunction with Calnexin and Calreticulin in the refolding of glucosidated proteins (34). Moreover, Δ*pdi1* cells showed a hypersensitivity response to DTT, thus confirming the canonical role of Pdi1 in *U. maydis* (Fig 5C and D).

To identify the causes behind the failure of Δ*pdi1* plant infections we first analyzed if the *pdi1* deletion causes growth alterations under axenic conditions. We found no differences in generation time (∼2.5 hours in rich media) (S2A Fig) or cell morphology (S2B Fig) between *wt* and Δ*pdi1* cells. Moreover, Δ*pdi1* did not show any defects in cell wall integrity or oxidative stress resistance (S3 Fig). Next, we examined if the mating capability between sexually compatible cells depends on Pdi1. This can be tested by analyzing dikaryon filament formation on charcoal plates, which are recognizable by the formation of white fuzzy colonies. We observed similar mating efficiency in FB1Δ*pdi1* and FB2Δ*pdi1* crosses compared to the *wt* strains (Fig 6A). Therefore, the pathogenic defects affecting Δ*pdi1* infections are probably not caused by failures in cellular growth or mating stages. Hence, the plant infection defects observed in *pdi1* mutant cells probably appear during interaction with the host. In fact, we have observed an increase in both Pdi1 protein levels upon *biz1* activation and *pdi1* transcription during the infection process, with a peak at three days post infection (Fig 7), in agreement with a high-throughput transcriptomic analysis of pathogenesis (35). These observations suggest that Pdi1 function is required during the plant infection process, possibly to ensure the correct folding of the increased quantity of glycosylated and secreted proteins that must be expressed for a successful infection to develop.

**Fig 6.**
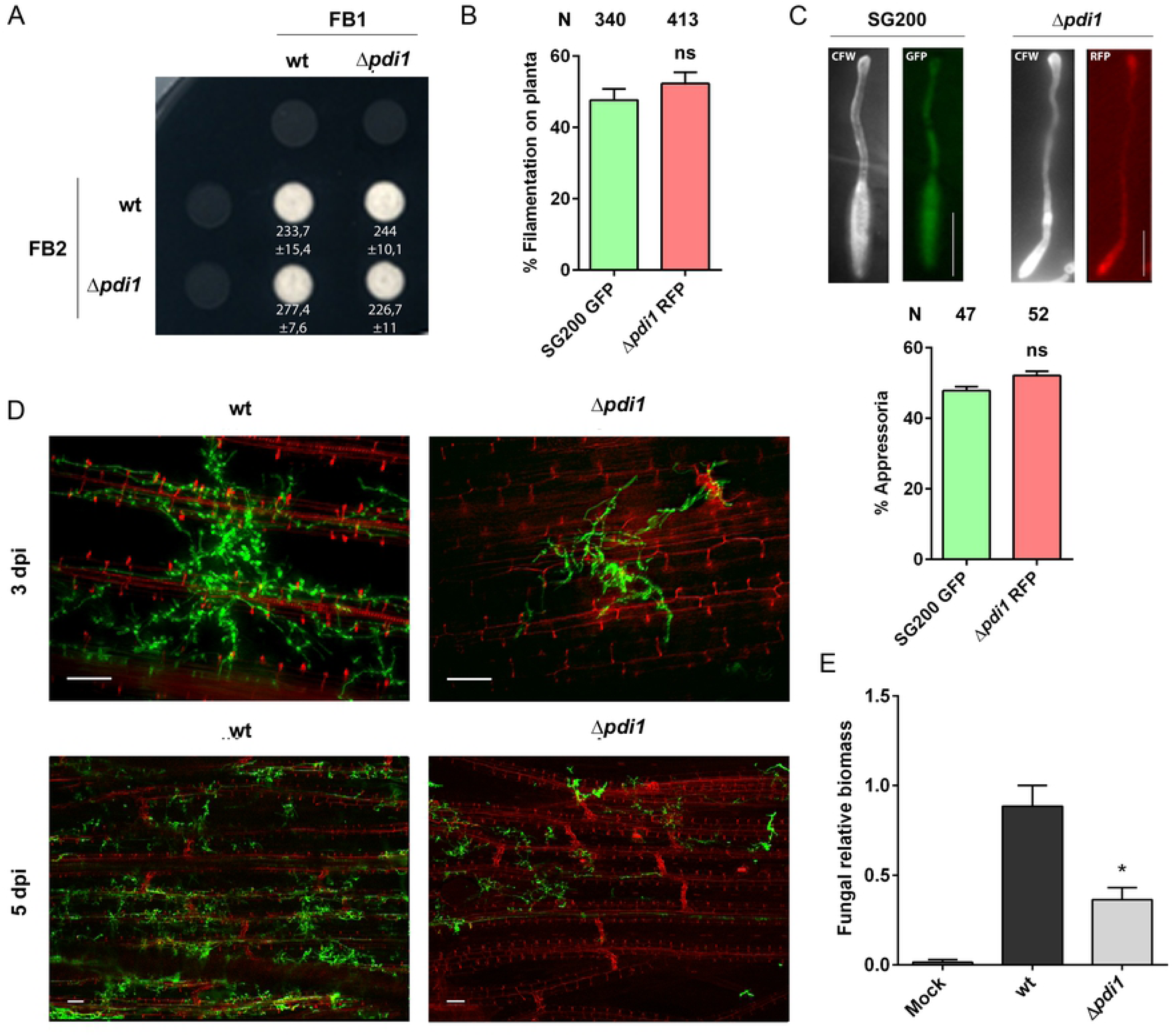
Pdi1 is essential for fungal growth within the maize plant. Mating assay between compatible *U. maydis* strains, FB1 and FB2 Δ*pdi1* strains were tested on PD-charcoal (PD-Ch) plates (A). Maize seedlings were co-infected with SG200 GFP and SG200 Δ*pdi1* RFP strains. After 20h, filament (B) and appressoria (C) formation were measured by scoring for GFP or RFP fluorescence. Total number of plants infected is indicated above each column. Scale bar represents 5 µm. (D) Maize leaves from 3 and 5 dpi plants infected with SG200 and SG200Δ*pdi1* were stained with propidium iodide (red) and *U. maydis* hyphae with WGA-AF-488 (green), and visualized by fluorescence microscopy, showing a decrease in the growth and branching capability of the *pdi1* mutant. Scale bar represents 50 µm. (E) Quantification of fungal biomass in planta at 3 dpi was performed by qPCR, measuring the constitutively-expressed *ppi1 U. maydis* gene normalized to the constitutively-expressed plant *gapdh* gene, confirming its defective proliferation. T-test statistical analysis was performed (* for p-value < 0.05).

**Fig 7.**
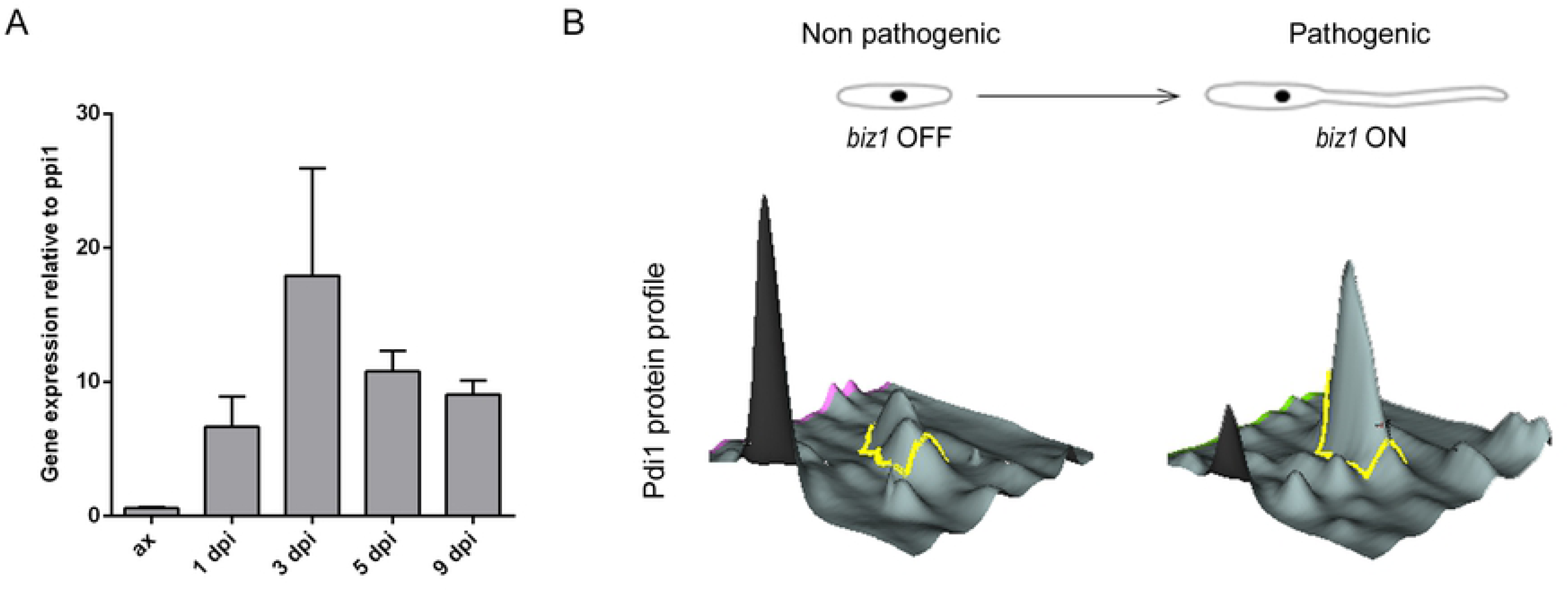
Pdi1 mRNA and protein expression profiles. (A) *pdi1* expression levels relative to *ppi1* during maize plant infection calculated by qRT-PCR from RNA isolated from plants infected with FB1 x FB2 at 1, 3, 5 and 9 days post infection and RNA from axenic culture. (B) Pdi1 protein expression profile before and after *biz1* activation obtained by DiGE analysis.

To determine which step of *U. maydis* pathogenic development is affected by the loss of Pdi1, we analyzed the behavior of Δ*pdi1* cells during plant cuticle pre-penetration stages. A physicochemical plant signal is recognized by the fungal cell, triggering filamentous growth and appressoria formation (18). In order to quantify both phenotypes, we generated the *pdi1* deletion in SG200 cells carrying RFP under the control of the constitutive *otef* promoter and co-inoculated maize plants with an equal mixture of SG200 P_otef_GFP and SG200 P_otef_RFP Δ*pdi1* cells. The amount of filaments and appressoria in *wt* and Δ*pdi1* cells was quantified using fluorescence microscopy ∼15 hours after infection. We found that the absence of Pdi1 did not alter filament formation capability (Fig 6B). Moreover, Δ*pdi1* filaments developed appressoria normally (Fig 6C). Thus, plant penetration stages do not require Pdi1, which suggests that deficient fungal expansion inside plant tissues might be behind the Δ*pdi1* infection defects.

To address the state of fungal colonization inside plant tissues we stained infected maize leaves 3 and 5 days after inoculation using WGA-Alexa and propidium iodide to visualize fungal hyphae and plant cells, respectively (see Methods). Following this approach, we observed a defective proliferation of Δ*pdi1* hyphae inside the plant (Fig 6D). To confirm these observations, we quantified fungal biomass in plant samples. We used the expression level of the *U. maydis ppi1* gene as a fungal marker (36). We found that the amount of fungal biomass 3 days post-infection decreased two-fold during Δ*pdi1* infection versus infection with a *wt* strain (Fig 6E). Thus, Pdi1 plays a key role in *U. maydis* colonization after plant penetration. Interestingly, this phenotype is reminiscent of defects caused by N-glycosylation defective cells (5) suggesting that altered Pdi1 function might be partly responsible for the defects associated with the loss of N-glycosylation.

### Loss of Pdi1 N-glycosylation mimics Δ*pdi1* virulence defects

An important remaining question is whether the loss of Pdi1 glycosylation mediated by Pmt4 and Gls1 leads to dysfunctional Pdi1 and a reduction in virulence similarly to Δ*pdi1* cells, or alternatively, a non-glycosylated Pdi1 form is still able to induce plant tumors and Pmt4 and Gls1 affect virulence elsewhere. To address this issue, putative O- and N-glycosylation sites were substituted with alanine (S4A, S24A, S340A, S344A, S346A, S350A, S477A and T486A) and glutamine (N36Q and N484Q) residues, respectively (Methods, Fig 8A). Pdi1 alleles harboring substitutions at all O-glycosylation sites (Pdi1^ΔO-gly^), all N-glycosylation sites (Pdi1^ΔN-gly^), all O- and N-glycosylation sites (Pdi1^ΔN,O-gly^) or the wild type allele were introduced in the exogenous *ip* locus of Δ*pdi1* cells under the control of the *otef* promoter and their ability to infect maize plants tested. We found that while Pdi1^ΔO-gly^ restored the virulence defects observed in Δ*pdi1* cells during infection similar to the *wt* allele, Pdi1^ΔN-gly^ and Pdi1^ΔN,O-gly^ forms failed to complement the lack of Pdi1 (Fig 8B). This strongly suggests that only the N-glycosylation of Pdi1 is relevant for its role in virulence. Similarly to *Δpdi1* cells, the virulence defects observed in Pdi1^ΔN-gly^, are due to a failure in the fungal expansion inside the plant tissues (Fig 8C). Remarkably, Pdi1^ΔN-gly^ could be detected by western blot discarding a possible degradation of Pdi1 in the absence of proper N-glycosylation (Fig. 8D). Consistent with defective Pdi1 function during plant infection when N-glycosylation target residues are removed, we observed that Pdi1^ΔN-gly^ does not complement the DTT sensitivity shown by the Pdi1 deletion mutant (Fig 8E).

**Fig 8.**
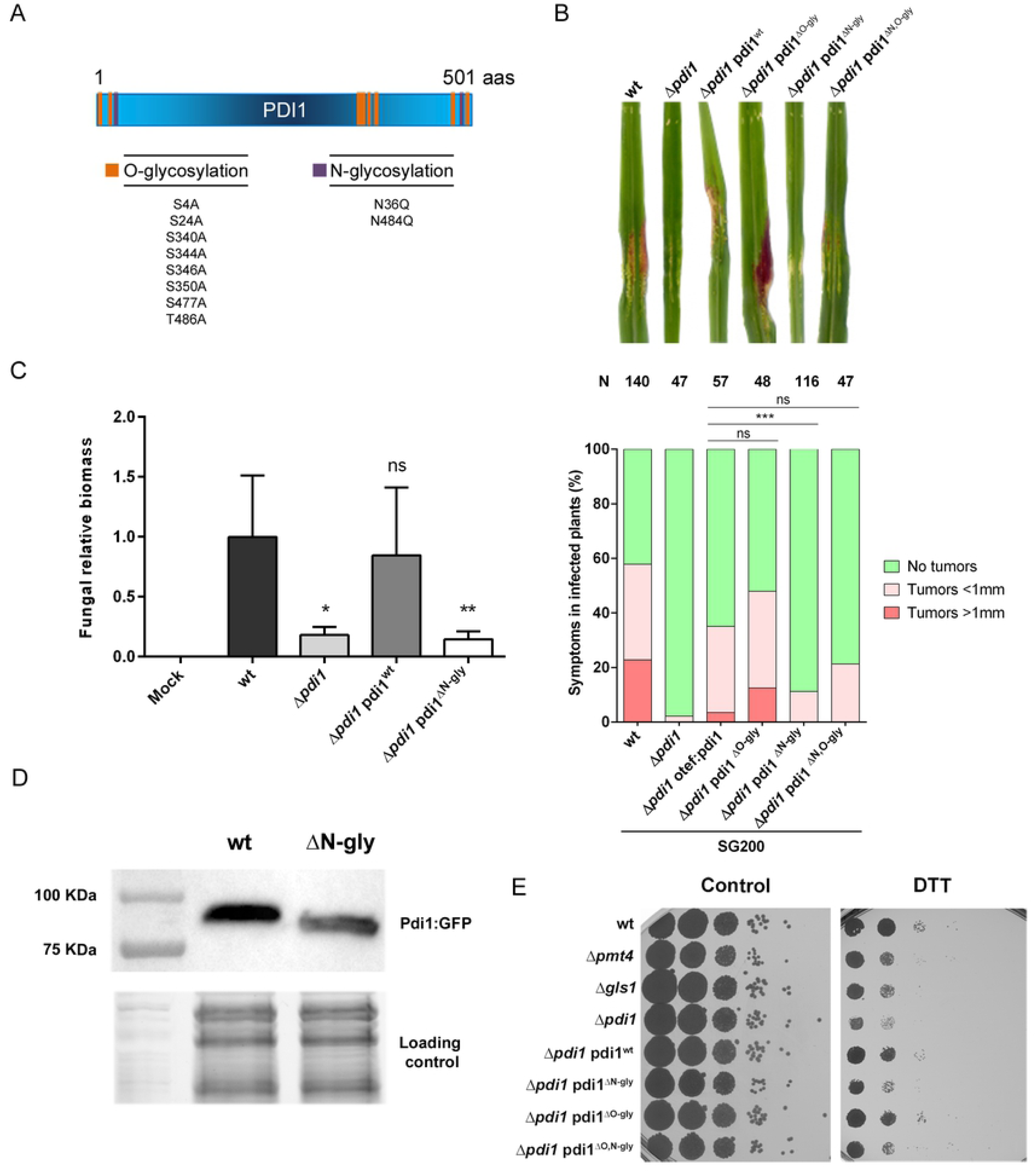
N-glycosylation of the protein disulfide isomerase Pdi1 ensures *U. maydis* virulence. (A) Pdi1 has two putative N-glycosylation sites and eight putative O-glycosylation sites. The main amino acids where the glycosylation tree is anchored, serine/threonine in O-glycosylation and asparagine in N-glycosylation, have been replaced by similar amino acids alanine and glutamine for O- and N-glycosylation, respectively. (B) The percentage of symptoms in maize plants infected with the indicated strains at 14 dpi. The total number of infected plants is indicated above each column. Mann-Whitney statistical test was performed (ns: not statistically significant; **** for p-value < 0.0001). Representative disease symptoms are shown above. (C) Quantification of fungal biomass in planta at 5 dpi was performed by qPCR, measuring the constitutively-expressed *ppi1 U. maydis* gene normalized to the constitutively-expressed plant *gapdh* gene. T-test statistical analysis was performed (ns for not statistically significant, * for p-value < 0.05, ** for p-value < 0.01). (D) Western blot showing Pdi1:GFP in SG200 and Pdi1_ΔN-glycosylation:GFP in SG200 Δ*pdi1*, with the image of the stain-free gel activation as a loading control. (E) Indicated strains were spotted onto CM plates supplemented with 2% D-glucose and 4 mM DTT. The mutation of N-glycosylation sites resulted on high DTT sensitivity.

### Δ*pdi1* and Δ*gls1* show similar secreted protein defects

Taking our results together with the established functions of Pdi1 in other model organisms lead us to propose a model whereby Gls1-mediated glycosylation of Pdi1 is required for correct folding and subsequent secretion of proteins activated by the virulence program. To explore this hypothesis, we compared the secretomes of *wt*, Δ*pdi1* and Δ*gls1* cells upon Biz1 induction (pathogenic filamentous growth) conditions. We found 6 secreted proteins showing differential electrophoretic mobility in *wt versus* Δ*pdi1* cells: Um06158 (probable glutaminase A), Um01829 (Afg1, previously identified in our first screening, see Fig 1), Um04503, an α-N-acetylgalactosaminidase, and Um01213, Um00027 and Um11462 (3 uncharacterized proteins) (see Fig 9). Remarkably, 4 out of these 6 proteins (Um06158, Um01829, Um04503 and Um00027) were also identified in the comparison between *wt* and Δ*gls1* secretomes suggesting that the effects caused by Δ*gls1* might be explained by deficient Pdi1 function. This indicates that there is a pool of secreted proteins that are glycosylated by Gls1 and their proper folding is dependent on Pdi1, which itself requires Gls-mediated glycosylation for function. The fact that the number of glycoproteins altered by the loss of *gls1* is higher than by *pdi1* is probably because a group of N-glycoproteins does not require Pdi1 for correct folding, secretion and function. This is consistent with the fact that Δ*gls1* causes more severe virulence defects than Δ*pdi1* (See model Fig 10).

**Fig 9.**
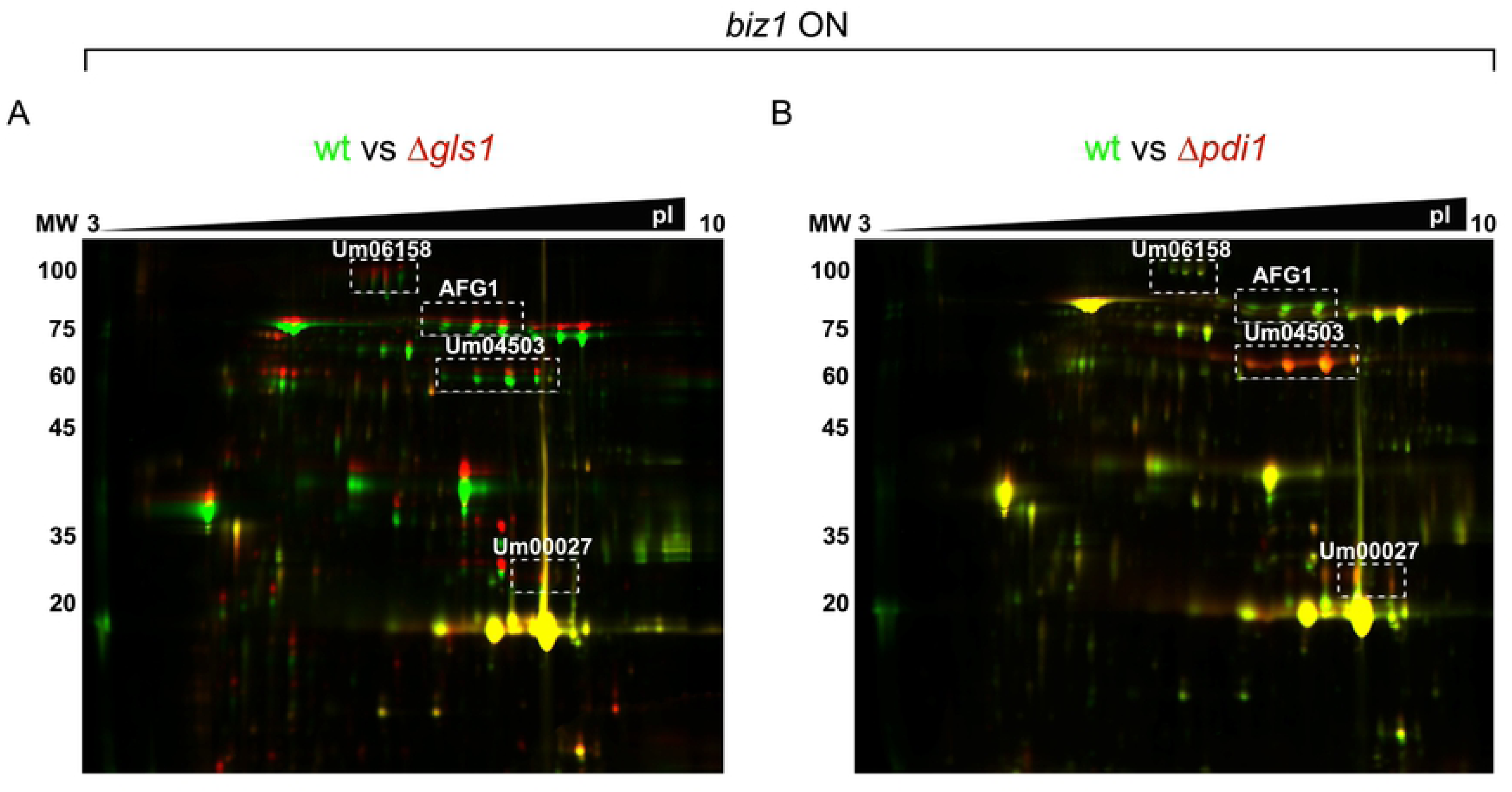
2D-DIGE expression analysis of Gls1- or Pdi1-dependent secreted glycoproteins. DIGE gel images showing protein changes between wild-type *biz1*^*crg*^ (tagged with Cy3 – green) and *gls1* (tagged with Cy5 – red) mutants (A) or between wild-type *biz1*^*crg*^ (tagged with Cy3 – green) and *pdi1* (tagged with Cy5 – red) mutants (B). Several glycoproteins depending on both Gls1 and Pdi1 could be identified. Three such proteins are indicated on the gels by dotted rectangles.

**Fig 10.**
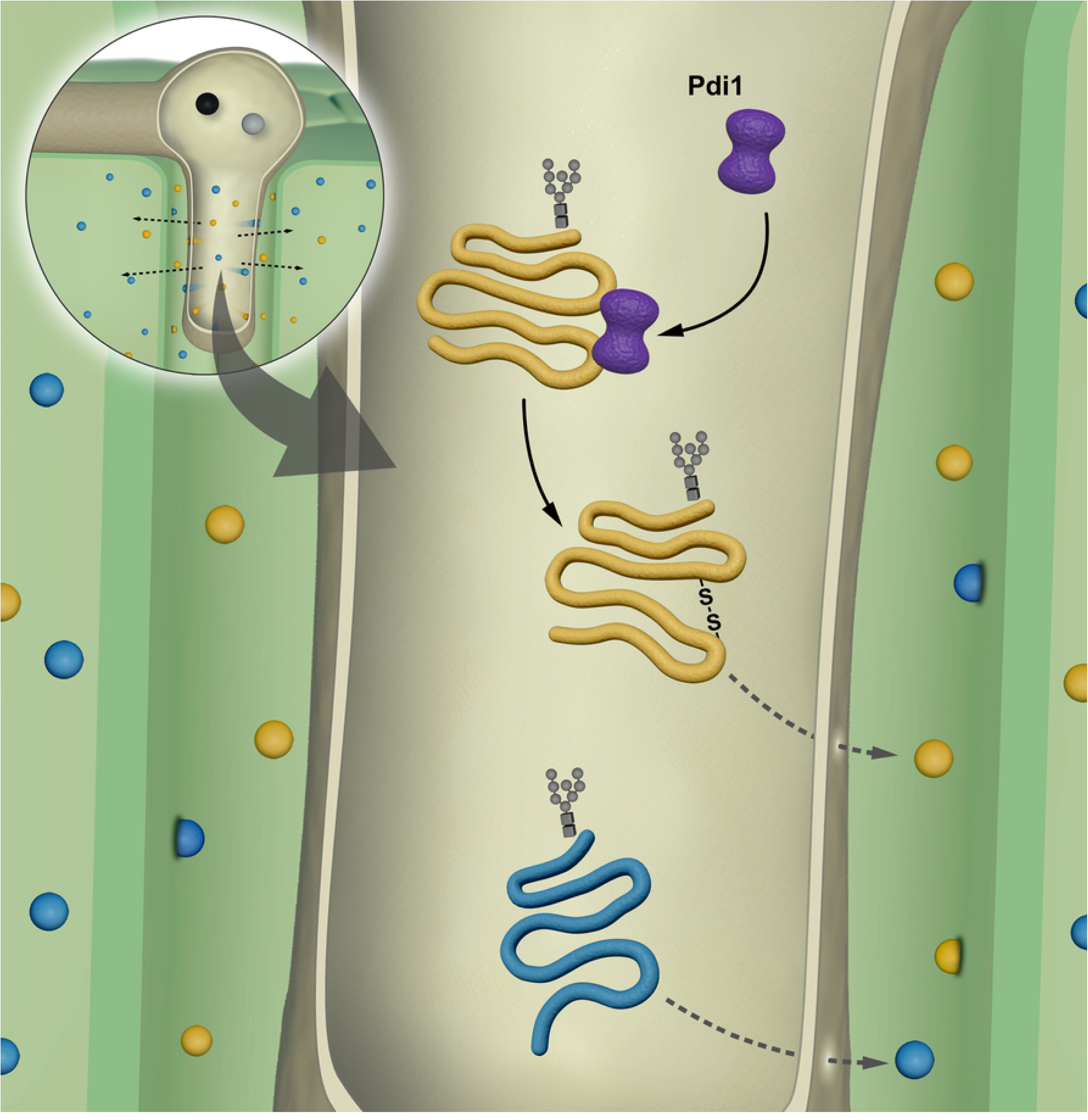
Model of Pdi1 function during infection. Pdi1 assists a subset glycoprotein in folding and disulfide bonds formation during effector secretion in the apoplast of maize plant during *U. maydis* infection.

## Discussion

In this work, we have demonstrated that the use of glycosylation mutants Δ*pmt4* (4) and Δ*gls1* (5) in a virulence program-specific 2D-DIGE protein analysis is a suitable and effective approach for identifying *Ustilago maydis* glycoproteins involved in plant infection. We identified 35 glycoproteins, and showed that 14 out of the 28 (50%) assayed are involved in virulence, analyzing single deletion mutants. Of these mutants, loss of the protein disulfide isomerase Pdi1 had the strongest effect on virulence, most likely due to protein folding defects and changes in functionality and/or secretion of proteins required for plant infection.

### An effective method for identifying new glycosylated fungal effectors

It is well established that protein glycosylation is essential for fungal infection. Mutants for specific glycosyltransferases or glycosidades are affected at different stages of the infection process in *Ustilago maydis* (4,5), other plant pathogens such as *Penicilium digitatum* (37), *Margnaporthe oryzae* (15,38) and *Botrytis cinerea* (39) as well as in animal pathogens such as *Candida albicans* (13,40) and *Cryptococcus neoformans* (6,41,42). Thus, our interest focused on determining which of the glycosylated proteins produced during the infection process were responsible for those defects. We postulated cell wall and secreted proteins as the main sources of novel virulence factors. Cell wall proteins form the pathogen’s most external layer while secreted factors have well established roles as effector proteins (43). Significantly, almost all cell wall and secreted factors are glycosylated (44,45). *U. maydis* putatively secretes 467 effectors that contribute to the establishment of biotrophy with maize during the infection process (19,22). In-silico tools such as SignalP (46), ApoplastP (47) or EffectorP (48) can predict which proteins are secreted and could be putative effectors. Although useful guides, such bioinformatic tools are not completely reliable since it has also been demonstrated that some fungal effectors are not secreted through the canonical secretion pathway involving N-terminal signal peptide degradation prior to translocation to the ER (49–52). For example, while around 90% of proteins secreted by *Fusarium graminearum* in axenic culture possess a signal peptide, only 56% of proteins secreted during the infection process have secretion signals (53). Different strategies such as RNA-seq and microarray, have been employed in *Ustilago maydis* in order to identify effectors whose transcription is induced during infection (54–57). Such studies highlight the huge changes in gene expression that occur during different stages of the infection process. Nevertheless, none of these approaches took into consideration the importance of protein modification, which are especially important for secreted proteins. For this reason, a proteomics approach can be an attractive alternative to identify proteins with important roles in the plant infection process. Hence, in this work we established a new approach to identify glycosylated effectors based on their electrophoretic mobility. Alterations to the glycosylation status of a protein in O- or N-glycosylation mutant strains, Δ*pmt4* or Δ*gls1* respectively, can be detected by bidimensional gel electrophoresis. This strategy allowed us to identify 27 glycoproteins from the cytosolic extract, 11 from the secreted protein extract and 6 from the cell wall extract, with some detected in multiple extracts. It is important to highlight that this approach identified previously characterized proteins involved in virulence as well as known effectors such as Afg1 and Afg3 (28), Pep4 (27) and Cmu1 (26). Among the 35 proteins identified, 4 are cell wall proteins that could be related to plant interaction during infection and 18 are located (Djamei, personal communication) and/or predicted to be located at the apoplastic region (S1 Table), which is consistent with effector protein function. Similar proteomic strategies based on glycosylation changes after the activation of virulence programs have been used to identify new fungal effectors in other pathogenic fungi. For example, the controlled induction of Mst11, a *Magnaporthe oryzae* MAP kinase essential for appressoria formation and plant infection (58) has been assayed in O-glycosylation mutants such as Δ*MoPmt2* (38) and Δ*MoPmt4* (59), as well as in N-glycosylation mutants such as Δ*alg3* (15) and Δ*Mogls2* (60).

Although it has been described that many single effector gene deletions do not affect plant infection (22), half of the glycoprotein deletions examined in a CL13 background showed a reduction in the number or size of tumors when maize plants were infected (Fig 4). In fact, 14 of the 28 single deletion mutants assayed showed a statistically significant decrease in the number of tumors. Interestingly, four of these deletions induced only very small (<1 mm) tumors. Ten of the most promising candidates were then deleted in the SG200 strain and tested for virulence in maize plants, showing that *Δpdi1* was the least virulent mutant. Single deletions for the other genes in this background caused little or no reduction in virulence. This observation may be explained by the fact that the CL13 strain does not possess the pheromone encoding gene *mfa2* (29) resulting in a reduced ability to develop filaments and produce tumors during maize plant infection. Thus, CL13 is more sensitive to the deletion of genes with minor but important contributions to the infection process.

Total disruption of the whole glycosylation process in *U. maydis* leads to avirulent phenotypes because of its inability to develop functional appressoria (Δ*pmt4*) or due to failure to spread inside the maize plant (Δ*gls1*) (4,5,61), however we did not find a single factor which resembles these phenotypes. The reason why this single factor was not identified might be because *U. maydis* instead uses a pool of glycosylated effectors that work cooperatively to perform the different steps involved in plant invasion and colonization.

### Pdi1 is a key factor supporting *U. maydis* pathogenic development

Of all the virulence-related glycoproteins identified here, we chose to further characterize Pdi1 as its deletion led to the strongest reduction in virulence. As we expected, Pdi1 localized to the ER and its deletion led to growth defects in axenic culture when an ER stress-inducing drug such as DTT was added to either solid or liquid media (Fig 5). These results support the idea that Pdi1 is a disulfide-isomerase protein that might assist the folding of newly synthetized proteins in the ER *via* the addition of disulfide bonds, which is critical during ER stress conditions (33). Many *U. maydis* effector proteins harbor cysteine-rich regions suitable for the formation of disulfide bonds, which may be necessary for their acquisition of active conformations (22,62,63). Thus, Pdi1’s role in establishing disulfide bonds in proteins involved in plant infection could be a major part of how the Δ*pdi1* mutation affects virulence. Moreover, both calreticulin (CRT) and calnexin (CNX), which bind glycosylated N-glycans in the ER, have a Pro-rich arm that may bind a protein disulfide isomerase (34), such as Pdi1. N-glycans may also play an important role in ER-associated degradation (ERAD) of proteins (34). Consequently, Pdi1 may be directly or indirectly involved in the selection of misfolded glycoproteins for dislocation into the cytosol for degradation by proteasomes, i.e., through N-glycans and mannosidase 1 (Mns1) and a second set of proteins called MnlI (mannosidase-like), Htm1 (homologous to mannosidase), or ER degradation-enhancing α-mannosidase-like protein (EDEM) (1,34,64,65).

We have demonstrated that the Pdi1 deletion strain can complete the initial steps of the infection process, appressoria formation and dikaryotic filament formation normally (Fig 6). Inside the maize plant, this mutant is able to form clamp-like cells and hyphae spread into maize cells although it cannot reach the vascular tissue. This growth defect was also observed by qPCR fungal biomass quantification, with more than a 50% reduction in fungal biomass versus wild-type (Fig 6). During *U. maydis* biotrophic development until tumor formation at 6 dpi, the fungus has to form the dikaryotic filament, develop appressoria, penetrate into the plant, modify plant metabolism to avoid activation of the plant’s immune system and ensure its proper propagation inside the host. In order to achieve all these tasks, *U. maydis* secretes several early effectors such as Pep1 (66,67), Pit2 (68), Stp1 (35) and See1 (69). As Δ*pdi1* did not show any defects in appressoria formation, dikaryotic filament formation or maize plant penetration, the reduction in fungal biomass observed in this mutant may be the result of a failure of the folding and secretion of proteins to the apoplast. Therefore, we propose three non-mutually exclusive mechanisms by which the loss of Pdi1 could affect fungal virulence: 1) a decrease or abolition in the secretion of effectors; 2) a reduction or elimination of the activity of the secreted effectors; 3) the presence of abnormally folded proteins that behave as new pathogen-associated molecular patterns (PAMPs) and activate the plant immune system. Work towards deciphering the molecular basis of Pdi1 action during plant infection and testing the above hypotheses is currently ongoing in our laboratory. Our results raise the possibility that mechanisms linking glycosylation and disulfide bond formation of fungal effectors are conserved and highlight the need to explore Pdi1’s role in other fungal pathogens.

## Abbreviations

ER: Endoplasmic Reticulum
GA: Golgi Apparatus
DIGE: Differential in Gel Electroforesis
HPLC: High Performance Liquid Chromatography
DTT: Dithiothreitol
MALDI-MS: Matrix-Assisted Laser Desorption/Ionization – Mass Spectrometry
ERAD: ER-Associated Degradation
dpi: days post-infection
PAMPs: Pathogen-Associated Molecular Patterns
CWP: Cell Wall Protein
IEF: Isoelectric Focusing

## Acknowledgements

We thank Antonio Pérez-Pulido for help with the *in silico* screening of glycosylation sites, Armin Djamei for sharing data about apoplastic proteins, and John R. Pearson for critical comments on the manuscript. We would like to thank the Genetics Department for their useful discussions and comments, Víctor Manuel Carranco for the graphic design in the figures and Sandra Romero for technical support. MMM was supported by P09-AGR-5241 Junta de Andalucía. IMS was awarded by BES-2014-069149 MINECO AEI/FEDER, UE, Spain. This work was supported by Junta de Andalucía P09-AGR-5241 grant and Spanish Government BIO2013–48858-P and BIO2016-80180-P grants from MINECO AEI/FEDER, UE and Ramon y Cajal program, RyC-2016-19659 to AFA.

## Materials and Methods

### Strains, plasmid and growth conditions

*Escherichia coli* DH5α and pGEM-T easy (Promega) and pJET1.2/blunt (ThermoFisher Scientific) were used for cloning purposes. Growth conditions for *E. coli* (70) and *U. maydis* (71) have been previously described. *U. maydis* strains used in this study are listed in S2 Table.

To induce the over-expression of transcription factor Biz1 (25), FB1Biz1^crg^ and its derivative mutants (FB1Biz1^crg^Δ*pmt4*, FB1Biz1^crg^Δ*gls1* and FB1Biz1^crg^Δ*pdi1*) cells were grown at 28°C in complete medium (CM) supplemented with D-glucose 25% (CMD), washed twice with water and grown in CM supplemented with arabinose 25% (CMA) at 28°C for 8 hours.

Pathogenicity assays were performed as described in (19). *U. maydis* cultures were grown at 28°C to exponential phase in liquid YEPSL (0.4% bactopeptone, 1% yeast extract and 0.4% saccharose) and concentrated to an OD_600_ of 3, washed twice in water and injected into 7 days old maize (*Zea mays*) seedlings (Early Golden Bantam). Disease symptoms were quantified 14 dpi. Statistical analyses were performed in GraphPad Prism 6 software.

ER stress assays were carried out with cultures grown at 28 °C to exponential phase in CMD and spotted at 0.4 OD_600_ onto CM plates supplemented with 4 mM DTT (iNtRON Biotechnology). Plates were incubated for 48 h at 28 °C. For ER stress assay in liquid culture, *U. maydis* cells were grown to exponential phase in YEPSL and diluted to 0.1 OD_600_ in YEPSL with 10 mM DTT. Cell growth at 28 °C with continuous shaking was analyzed over 24 h using a Spark 10M (Tecan) fluorescence microplate reader.

Cell wall integrity and oxidative stress assays were carried out with cultures grown at 28 °C to exponential phase in CMD and spotted at 0.4 OD_600_ in CM plates supplemented with calcofluor white 40 µg/ml (Sigma-Aldrich), Congo Red 50 µg/ml (Sigma-Aldrich), Tunicamycin 1 µg/ml (Sigma-Aldrich), Sorbitol 1M (Sigma-Aldrich), 2% DMSO (Sigma-Aldrich), H_2_O_2_ 1.5 mM (Sigma-Aldrich) and NaCl 1M (Sigma-Aldrich). Plates were incubated for 48 h at 28 °C.

For mating and filamentation assays, cells were grown in liquid YEPSL until exponential phase, washed twice with sterile bidistilled water, spotted onto PD-charcoal plates and grown for 24-48 hours at 25-28°C.

### DNA and RNA procedures

Molecular biology techniques were used as described in (70). *U. maydis* DNA isolation and transformation were carried out following the protocol described in (72).

To generate deletion mutants, 1 kb fragments of the 5’ and 3’ flanks of the gene of interest (*goi*) ORF were generated by PCR using Phusion High Fidelity DNA polymerase (New England Biolabs) or Q5 High-Fidelity DNA polymerase (New England Biolabs) and *U. maydis* FB1 genomic DNA, using the primers *goi*KO5-1 and *goi*KO5-2 (containing a *SfiI* restriction site) to amplify the 5’ flank and *goi*KO3-1 (containing a *SfiI* restriction site) and *goi*KO3-2 to amplify the 3’ flank (S3 Table). These fragments were digested with *SfiI* and ligated with the 1.9-kb *SfiI* carboxin, 1.4-kb *SfiI* noursethricin (*ClonNAT*), 2-kb *SfiI* geneticin or 1.9-kb *SfiI* hygromicin resistance cassettes (73). Constructs were cloned into pGEM-T easy (Promega) or pJET1.2/blunt (ThermoFisher Scientific) plasmids and amplified by PCR using the primers *goi*KO5-1/*goi*KO3-2, prior to their transformation in *U. maydis*.

To generate SG200 2xRFPΔ*pdi1* used for filament quantification into the maize plant, pGEM-TΔ*pdi1:cbx* plasmid was digested with *SfiI* to excise the carboxin resistance cassette and ligated to a hygromicin resistance cassette isolated from pMF1h (73) with *SfiI*/*BsaI* double digestion, leading to pGEM-TΔ*pdi1:hyg*. This construct was amplified by PCR with Phusion DNA polymerase using primers pdi1KO5-1/pdi1KO3-2 and integrated into the SG200 2xRFP strain (74).

For the SG200 3xGFP strain used for filament quantification in planta, the pOG plasmid (74), containing 3xGFP under the control of the *otef* promotor and a hygromicin resistance cassette, was digested with *PsiI* and integrated into the SG200 strain. Finally, Δ*pdi1:cbx* was amplified by PCR using primers pdi1KO5-1/pdi1KO3-2 and integrated into the SG200 3xGFP strain.

To perform *pdi1* deletion complementation assays, we generated the SG200Δ*pdi1*P_otef_*pdi1* strain. To this end, the eGFP fragment in p123 (75) was substituted with the *pdi1* ORF. The *pdi1* ORF was amplified by PCR using Phusion HF DNA polymerase (New England Biolabs) with primers pdi1StartXmaI and pdi1StopNotI, containing XmaI and NotI restriction sites, respectively. The PCR product was digested with XmaI and NotI, purified and cloned into the p123 plasmid digested with both restriction enzymes, creating p123-*pdi1*. Finally, p123-*pdi1* was linearized with *SspI* and integrated into the SG200Δ*pdi1 ip* locus by homologous recombination.

To create an O-glycosylation deficient *pdi1* mutant, serine or threonine sites predicted by NetOgly 4.0 tool were replaced by site-direct mutagenesis using GenScript (Piscataway, USA), generating the plasmid pUC57-*pdi1*mutO-gly, with *Sma1* and *NotI* restriction sites. The *pdi1*mutO-gly construct was reintroduced into p123 as described above, creating the p123-*pdi1*mutO-gly vector. This plasmid was linearized with *SspI* and integrated into the SG200Δ*pdi1 ip* locus by homologous recombination, generating the SG200Δ*pdi1pdi*^Δ*O-gly*^ strain.

To generate an N-glycosylation mutant version of *pdi1*, asparagine 36 and 484 were replaced by glutamine in N-glycosylation sites through PCR using the Gibson Assembly^®^ Cloning Kit (New England BioLabs, Frankfurt, Germany) with primers PDI1Asn36mut-1 and PDI1Asn36mut-2 for the mutation N36Q and primers PDI1Asn484mut-1 and PDI1Asn484mut-2 for the mutation N484Q, using p123-*pdi1* as template, generating the plasmid p123-*pdi1*mutN-gly. This construction was integrated into the SG200Δ*pdi1 ip* locus (as described above), leading to SG200Δ*pdi1pdi*^Δ*N-gly*^ strain.

To generate the O- and N-glycosylation *pdi1* mutant, N36 and N484 was replaced by glutamine using PCR and the Gibson Assembly Cloning Kit, with primers PDI1Asn36mut-1/PDI1Asn36mut-2 for the N36Q mutation and PDI1Asn484mutThr486mut-1/PDI1Asn484mutThr486mut-2 for the N484Q mutation, and the plasmid p123-*pdi1*mutO-gly as template, reintroduced into p123 to generate p123-*pdi1*mutN,O-gly. This construct was linearized with *SspI* and reintegrated into the SG200Δ*pdi1 ip* locus.

All *pdi1* mutants for N- and/or O-glycosylation were confirmed by sequencing, using primers PDI1SeqI and PDI1SeqII.

For GFP tagging of *pdi1*, we generated plasmid pDL51-*pdi1*. This plasmid is a p123 derivative (16) where *pdi1* ORFs were PCR amplified using Phusion DNA polymerase with primers pdi1ORF5/pdi1ORF3, was cloned in frame with the GFP present in the plasmid, under the control of the constitutive expressed *otef* promotor. To achieve this, pDL51 was linearized with *SfiI* digestion and ligated with the *pdi1* ORF digested with *SfiI*. For GFP tagging of the N-glycosylation mutant version of PDI1, *pdi1*mutN-gly was amplified by PCR with primers pdi1StartXmaI and pdi1StopNcoI containing XmaI and NcoI restriction sites, respectively. The PCR product was digested with XmaI and NcoI, purified and cloned into the p123GFP plasmid digested with both restriction enzymes, to generate an in frame C-terminal *pdi1*mutN-gly GFP fusion. Finally, p123-*pdi1*mutN-glyGFP was linearized with *SspI* and integrated into the SG200Δ*pdi1 ip* locus by homologous recombination.

For fungal biomass quantification, 3 and 5 dpi maize leaves were ground in liquid nitrogen and total DNA was isolated with DNeasy Plant Mini kit (Qiagen) according to manufacturer’s instructions. Fungal biomass was then quantified by Real-Time PCR with an ABIPRISM 7000 Sequence Detection System (Applied Biosystems) using the Power SYBR Green PCR Master Mix (ThermoFisher Scientific) according to the manufacturer’s protocol, measuring the constitutively-expressed fungal *ppi1* gene, using primers RT-PPI1-5/RT-PPI1-3, and and constitutively-expressed plant *gapdh*, using primers Gapdh-F/Gapdh-R for normalization purposes. 100 ng of total DNA was used a template for each reaction.

For Pdi1 expression analysis, RNA from FB1 axenic culture and maize plants infected with FB1 and FB2 was isolated at 1, 3, 5 and 9 dpi, ground in liquid nitrogen using a pestle and mortar and purified using TRIzol reagent (Thermo Fisher Scientific) and the Direct-zol™ RNA Miniprep Plus kit (Zymo Research), following the manufacturer’s instructions. cDNA was synthetized using RevertAid H Minus First Strand cDNA Synthesis kit (Thermo Fisher Scientific) following the manufacturer’s instructions. Pdi1 expression levels were quantified by qRT-PCR using a Real-Time CFX Connect (Biorad) and SYBR® Premix Ex Taq™ II (Tli RNase H Plus) (Takara) according to the manufacturer’s protocol, measuring *pdi1* expression gene, using primers pdi1 RT-fwd/pdi1 RT-rev, and *ppi1* as a constitutively-expressed control, using primers RT-PPI1-5/RT-PPI1-3.

### Proteomic analysis

To detect pathogenesis-related changes to the cytosolic proteome caused by the loss of *pmt4*, we isolated O-glycosylated proteins from FB1Biz1^crg^ and FB1Biz1^crg^Δ*pmt4* strains. FB1 and FB1Δ*pmt4* strains were used as controls. Cells from each strain were grown in 1 L of CMD to an OD_600_=0.5-0.8, washed twice with sterile bidistilled water and grown for 8 hours in 1 L of CMA. After filamentation induction, samples were harvested by centrifugation, washed twice in ice-cold 20 mM Tris-HCl buffer (pH 8.8) and frozen at −80 °C. Cells were then resuspended in lysis buffer (20 mM Tris-HCl, 0.5 M NaCl, pH 7.4) containing protease inhibitor cocktail (cOmplete Tablets, EDTA-free, Roche) after which cell lysis was performed using glass beads (Sigma) in a FastPrep®-24 homogeniser (MP Biomedicals), with power set to 6.0 using 3 x 45” pulses at maximum speed with 5 min rest between cycles. After cell lysis, tubes were drilled with a needle and put into a new tube and samples were recovered by centrifugation at 4000 rpm for 1 min. Subsequently, samples were centrifuged at 14000 rpm for 30 min at 4°C and the supernatant was collected. One volume of 2x binding buffer (20 mM Tris-HCl, 0.5 M NaCl, 2 mM MnCl_2_, 2 mM CaCl_2_, pH 7.4) was added and concentrated using Amicon Ultra-15 3 kDa centricon (Merk Millipore) and O-glycosylated proteins isolated by affinity chromatography using Concavaline A columns (HiTrap Con A 4B, GE Healthcare) in AKTA FPLC (GE Healthcare), and eluted with a buffer containing 20 mM Tris-HCl, 0.5 M NaCl and 0.5 M methyl-α-D-manopyranoside (pH 7.4).

To detect pathogenesis-related changes to the cytosolic proteome caused by the loss of *gls1*, FB1Biz1^crg^ and FB1Biz1^crg^Δ*gls1* cells were grown in 1 L of CMD to an OD_600_=0.5-0.8, washed twice with sterile distilled water and grown for 8 hours in 1 L of CMA. FB1 and FB1Δ*gls1*, used as controls, were grown in 2 L of CMD to an OD_600_=0.5-0.8, washed twice with sterile distilled water and grown during 8 hours in 2 L of CMA. Samples were harvested by centrifugation, washed twice in ice-cold 20 mM Tris-HCl buffer (pH 8.8) and frozen by dropping samples in liquid nitrogen. Cells were lysed with 3-4 cycles in 6870 Freezer/Mill (SPEX, SamplePrep) according to the manufacturer’s instructions. Samples were resuspended in binding buffer (20 mM Tris-HCl, 0.5 M NaCl, 2 mM MnCl_2_, 2 mM CaCl_2_, pH 7.4) containing protease inhibitor cocktail (cOmplete Tablets, EDTA-free, Roche) and centrifuged at 14000 rpm for 30 min at 4°C. Supernatant was filtered and N-glycosylated proteins isolated by affinity chromatography using Concavaline A columns (HiTrap Con A 4B, GE Healthcare) in AKTA FPLC (GE Healthcare), and eluted with a buffer containing 20 mM Tris-HCl, 0.5 M NaCl and 0.5 M methyl-α-O-glucopyranoside (pH 7.4).

For differential analysis of secreted O- or N-glycoproteins in Δ*pmt4* or Δ*gls1* mutants, FB1Biz1^crg^ and FB1Biz1^crg^Δ*pmt4* or FB1Biz1^crg^Δ*gls1* cells were grown in 1 L of CMD to an OD_600_=0.5-0.8, washed twice with sterile bidistilled water and grown for 8 hours in 1 L of CMA. After filamentation induction, cells were centrifuged, the supernatant filtered and secreted glycosylated proteins precipitated with deoxycholate acid and TCA.

Cell wall O- or N-glycoproteins in Δ*pmt4* or Δ*gls1* mutants were purified following the protocol described in (76), with some modifications. FB1Biz1^crg^ and FB1Biz1^crg^Δ*pmt4* or FB1Biz1^crg^Δ*gls1* cells were grown in 1 L of CMD to an OD_600_=0.5-0.8, washed twice with sterile bidistilled water and grown for 8 hours in 1 L of CMA. After filamentation induction, samples were harvested by centrifugation, washed four times in ice-cold 10 mM Tris-HCl buffer (pH 7.4) and frozen at −80 °C. Cells were then resuspended in 10 mM Tris-HCl buffer (pH 7.4) containing protease inhibitor cocktail (cOmplete Tablets, EDTA-free, Roche) after which cell lysis was performed with glass beads (Sigma) in a FastPrep®-24 homogeniser (MP Biomedicals), with power set to 6.5 using 6 x 60” pulses at maximum speed with 3 minutes rest between cycles. Samples were recovered by centrifugation at 3000 x g for 10 min at 4°C, after drilling the bottom of the tube with a needle and placing into a new tube. The pellet (cell wall) was collected and washed sequentially with ice-cold sterile bidistilled water, 5% NaCl with protease inhibitor, 2% NaCl with protease inhibitor and 1% NaCl with protease inhibitor. Washing steps were repeated three more times and cell wall proteins (CWPs) purified after 10 min at 100 °C in extraction buffer (50 mM Tris-HCl pH 8.0, 100 mM EDTA, 2% SDS, 10 mM DTT) followed by 15 min centrifugation at 35000 x g. CWPs were then collected from the supernatant.

To analyze the common target between Gls1 and Pdi1, secreted proteins from FB1Biz1^crg^ and FB1Biz1^crg^Δ*gls1* or FB1Biz1^crg^Δ*pdi1* were precipitated with deoxycholate acid and TCA following the protocol described above for secreted O- and N-glycoproteins.

Isolated proteins for each fraction and genotype were then precipitated with 2D Clean-Up kit (GE Healthcare), and dissolved in TS buffer (7M urea, 2M thiourea, and 4% CHAPS) containing 30 mM Tris-HCl (pH 9.5). Protein concentration was determined using RC DC Protein Assay (Bio-Rad).

Fifty micrograms of protein extract from each sample were labeled with Cy3- or Cy5 Dye (GE Healthcare). Labeling was performed reciprocally so that each sample was separately labeled with Cy3 and Cy5 to account for any preferential protein labeling by the CyDyes. Twenty-five micrograms of each sample were labeled with Cy2-Dye and pooled together as an internal standard. Labeling was performed according to the manufacturer’s instructions. Different IEF procedures were performed. For cytosolic O-glycoproteins and CWPs, IEF was performed in 24 cm 4–7 NL Immobiline DryStrip (GE Healthcare) in an IPGphor unit (GE Healthcare) at 20 °C as follows: rehydratation for 12 h, 500 V for 1 h, linear gradient from 500 to 1000 V for 1 h, linear gradient from 1000 to 8000 V for 3 h and 8000 V to 60000 total Vhr for 5.5 h. The same IEF procedure using 24 cm 3-10 NL DryStrips was performed for secreted proteins. For cytosolic N-glycoproteins, IEF was performed using 24 cm 3–7 NL Immobiline DryStrip (GE Healthcare) in an IPGphor unit (GE Healthcare) at 20 °C as follows: rehydration for 12 h, 500 V for 1.5 h, linear gradient from 500 to 1000 V for 7 h, linear gradient from 1000 to 8000 V for 8 h and 8000 V to 60000 total Vhr for 5.5 h. After IEF, strips were incubated for 15 min at room temperature in a shaker in equilibration buffer (50 mM Tris-HCl [pH 8.5], glycerol 30% v/v, 6M urea, 2% w/v SDS) with 10 mg/mL DTT, followed by another 15 min incubation in equilibration buffer with 25 mg/mL iodoacetamide and loaded onto 10% acrylamide gels. Second dimension separation was performed using an EttanDalt Six Electrophoresis Unit (GE Healthcare).

After electrophoresis, gels were imaged using a Typhoon-9400 scanner (GE Healthcare) with a 100 µm resolution using appropriate emission and excitation wavelengths, photomultiplier sensitivity and filters for each of the Cy2, Cy3, and Cy5 dyes.

Relative protein spot quantification across experimental conditions was performed using DeCyder v7.0 software and multivariate statistical module EDA v7.0 (Extended Data Analysis, GE Healthcare) as follows. First, the Batch Processor module detected spots across the three gel images (two experimental samples and an internal standard) and generates differential in-gel analysis images with information about spot abundance in each image with values expressed as ratios. After spot detection, the biological variation analysis module utilizes differential in-gel analysis images to match protein spots across all gels, using the internal standard for gel-to-gel matching. Statistical analysis was then carried out to determine protein expression changes. Spot changes with a p-value lower than 0.01 calculated using Student’s t-test with the multiple testing assessed using the false discovery rate were considered significant. Multivariate analysis was performed by principal component analysis using the algorithm included in the EDA module of the DeCyder v7.0 software based on spots matched across all gels.

Protein identification was performed at the Proteomics Unit of the Pablo de Olavide University (Seville, Spain) and at the Proteomics service of Parque Científico de Madrid (Madrid, Spain). Spots were excised from the gels manually and transferred to 1.5 mL tubes. Sample digestion and MALDI-MS and MS/MS database searches were done by the Proteomics Units mentioned above.

### Western Blot analyses

For Western Blot analyses, cells were grown in YEPSL to an OD_600_=0.6-0.8 and were then collected by centrifugation at 4500 rpm for 5 min, washed twice with 20 mM Tris-HCl pH 8.8 and pellets were resuspended in lysis buffer (20 mM Tris-HCl, 0.5 M NaCL, pH 7.4) with protease inhibitor cocktail (cOmplete Tablets, EDTA-free, Roche). Samples lysis was performed with glass beads (Sigma) in a FastPrep®-24 homogeniser (MP Biomedicals), with power set to 6.5 using 6 x 60” pulses at maximum speed with 4 minutes rest between cycles. After cell lysis, tubes were drilled with a needle and put into a new tube and samples were recovered by centrifugation at 4000 rpm for 1 min. Subsequently, samples were centrifuged at 14000 rpm for 30 min at 4°C and the supernatant was collected. Protein concentration was measured using the RC DC Protein Assay kit (Bio-Rad). 60 μg of protein extract for each strain analyzed was loaded into a 10% TGX Stain-Free™ FastCast™ Acrylamide gel (Bio-Rad). Separated proteins were transferred onto a nitrocellulose membrane using the Trans-Blot® Turbo™ transfer system (Bio-Rad). The membrane was incubated with mouse polyclonal anti-GFP antibody (Roche) (1:1000). As secondary antibody, anti-mouse IgG-horseradish peroxidase conjugated antibody (1:5000; Sigma Aldrich) was used. Immunoreactive bands were developed by SuperSignal™ West Femto Maximum Sensitivity substrate (ThermoFisher Scientific). Image gel and membrane acquisition was carried out with ChemiDoc XRS (Bio-Rad).

### Microscopy

To corroborate ER Pdi1 localization, experimental SG200*pdi1:gfp* and ER localization control SG200*cal*^*s*^*:mrfp:HDEL* cells (77) were visualized using a spinning-disk confocal microscope (IX-81, Olympus; CoolSNAP HQ2 camera, Plan Apochromat 100×, 1.4 NA objective, Roper Scientific).

For DNA content visualization, cells were stained with DAPI and observed by Differential Interference Contrast (DIC) and fluorescence microscopy using a DeltaVision microscopy system comprising an Olympus IX71 microscope and CoolSnap HQ camera.

For *in planta* quantification of filament and appressoria formation in co-infection experiments with *U. maydis* GFP and RFP labeled strains, 20 dpi leaf samples were stained with calcofluor white (Sigma-Aldrich) to visualize fungal material and then checked for GFP or RFP fluorescence.

To analyze the *U. maydis* progression inside the maize plant, leaf samples from 3 and 5 dpi infected plants were distained with ethanol, treated 4h at 60 °C with 10% KOH, washed in phosphate buffer and then stained with propidium iodide (PI) to visualize plant tissues in red and wheat germ agglutinin (WGA)/AF488 to visualize the fungus in green. Samples were examined using a Leica SPE (DM2500) confocal microscope. Image processing was carried out using Adobe Photoshop CS5 and ImageJ.

## Supporting information

**S1 Fig. Um10156 is similar to *Saccharomyces cerevisiae* Pdi1.** Alignment of Pdi1 sequence from *S. cerevisiae* and Um10156 from *U. maydis* using T-Coffee Server and BoxShade Server. Thioredoxin domains are indicated by red lines.

**S2 Fig. Loss of *pdi1* has no effect on the non-pathogenic stage.** (A) Growth rate of SG200 and SG200 Δ*pdi1* in liquid rich media YEPSL. (B) SG200 and Δ*pdi1* cells observed by DIC microscopy don’t show any defect in size and morphology.

**S3 Fig. N-glycosylation mutation in Pdi1 does not affect cell wall integrity nor oxidative stress.** ER stress, cell wall integrity and oxidative assays were performed in CM plates supplemented with 2% D-glucose and calcofluor white (CFW) 40 µg/ml, Congo Red 50 µg/ml, Tunicamycin 1 µg/ml, Sorbitol 1M, 2% DMSO, H_2_O_2_ 1.5 mM and NaCl 1M.

**S1 Table. Identified glycoproteins in this study.** List of glycoproteins identified through the tree different extracts indicating in which extract were identified, their apoplastic secretion prediction and their gene deletion phenotypes after maize plant infection.

**S2 Table. List of strains used in this study.**

**S3 Table. List of primers used in this study.**

